# Closed ecosystems extract energy through self-organized nutrient cycles

**DOI:** 10.1101/2023.05.30.542875

**Authors:** Akshit Goyal, Avi I. Flamholz, Alexander P. Petroff, Arvind Murugan

**Affiliations:** Department of Physics, Massachusetts Insitute of Technology, Cambridge, MA 02139; Division of Biology and Biological Engineering, California Institute of Technology, Pasadena, CA 91125; Department of Physics, Clark University, Worcester, MA 01610; Department of Physics, University of Chicago, Chicago, IL 60637

## Abstract

Our planet is a self-sustaining ecosystem powered by light energy from the sun, but roughly closed to matter. Many ecosystems on Earth are also approximately closed to matter and recycle nutrients by self-organizing stable nutrient cycles, e.g., microbial mats, lakes, open ocean gyres. However, existing ecological models do not exhibit the self-organization and dynamical stability widely observed in such planetary-scale ecosystems. Here, we advance a new conceptual model that explains the self-organization, stability and emergent features of closed microbial ecosystems. Our model incorporates the bioenergetics of metabolism into an ecological framework. By studying this model, we uncover a crucial thermodynamic feedback loop that enables metabolically diverse communities to almost always stabilize nutrient cycles. Surprisingly, highly diverse communities self-organize to extract *≈*10% of the maximum extractable energy, or *≈*100 fold more than randomized communities. Further, with increasing diversity, distinct ecosystems show strongly correlated fluxes through nutrient cycles. However, as the driving force from light increases, the fluxes of nutrient cycles become more variable and species-dependent. Our results highlight that self-organization promotes the efficiency and stability of complex ecosystems at extracting energy from the environment, even in the absence of any centralized coordination.

---

The Earth surface is replete with ecosystems that are quasi-closed to material exchange, but open to light energy, e.g., lakes, microbial mats, and open ocean gyres [1–3]. Indeed nearly the entirety of Earth’s fossil record before plants and animals is composed of stromatolites—the mineral residues left over millennia from the activities of stratified microbial communities [4]. Winogradsky columns, a classic and key microbial experiment, are also examples of materially closed ecosystems [5–7]; these light-fueled closed columns seeded with mud show self-organization of nutrient cycles — of C, N, S, O, P — and are robust enough to use in undergraduate curricula. The Earth’s surface is itself a roughly materially closed ecosystem, with very little leakage to and from the mantle and space [8, 9]. All these closed ecosystems are self-organized and remarkably stable to perturbations; in fact, it is thought that being quasi-closed enables them to be extremely productive (high rates of nutrient (re)cycling), while maintaining a quasi-static non-growing state [10, 11]. However, we do not understand the principles that dictate when and why robust self-organized nutrient cycles emerge in either natural or synthetic ecosystems.

The central challenge of sustaining a closed ecosystem is the need to simultaneously solve several bioenergetic constraints without any central coordinator. Instead, closed ecosystems must self-organize via feedback mechanisms. In a materially closed system, light energy is the only external input, and can only be captured by recycling matter [12–15]. While directly accessible only to photosynthetic organisms, the energy in light must percolate through the ecosystem to sustain all organisms that perform different steps in recycling matter, i.e., different arms of nutrient cycles [16–20]. At the same time, the energy content of different nutrients depends on the recycling of metabolic products by other species. This suggests that the crucial feedbacks in closed ecosystems are thermodynamic in nature.

Despite these constraints, ecosystems stably re-establish nutrient cycles after major perturbations [21] — even after some of the largest perturbations observed in the history of life on Earth, e.g., the oxygenation of the atmosphere [22], ice ages [23, 24], and bolide impacts and mass extinctions of plants and animals [25]. Is the emergence, stability, and resilience of these self-organized non-equilibrium systems surprising? Further, the rules of self-organization are locally greedy — each species grows if it can extract sufficient energy for itself. How efficient should we expect ecosystems to be at extracting energy from light, given that they self-organize through such locally greedy rules?

Prevailing ecological models like consumer-resource models are successful at describing open ecosystems but cannot correctly capture the thermodynamic constraints and feed-backs key to closed ecosystems [26–28] Thus, in these models, closed ecosystems exponentially dwindle and collapse. Having theoretical models of stabilizing feedbacks in closed ecosystems is vital to understanding their stability to perturbations, efficiency at extracting energy, as well as conserved features across distinct ecosystems that emerge from the underlying constraints.

Here, we propose and study a theoretical framework of closed ecosystems using a redox framework, which incorporates the bioenergetics, conservation laws and thermodynamics of metabolism. These key features are missing from consumer-resource models, but become important in closed ecosystems, where the need to recycle products at balanced rates imposes tight thermodynamic constraints. We studied the emergence and stability of multiple nutrient cycles in a closed setting. By simulating our model, we found that once enough species are added, ecosystems almost always self-organize to a state where they can spontaneously recycle multiple nutrients, resulting in energy extraction. Even though distinct ecosystems contain different organisms, the fluxes of their nutrient cycles are in a very similar configuration (convergent). Further, the fluxes of self-organized nutrient cycles are stable to perturbations in species abundances and rapidly recover via thermodynamic feedback. Remarkably, in all these cases, ecosystems are very efficient at extracting energy. The energy extracted from spontaneous cycling is *≈*10% of the maximum extractable energy, or *≈*100 fold more than randomized communities. These results advance our understanding of how several coupled nutrient cycles spontaneously emerge and stabilize as a result of many interacting components. They also highlight the efficiency with which ecosystems extract energy from an external source like light. Finally, our work establishes closed ecosystems as paradigmatic examples of systems that self-organize to determine their displacement from equilibrium, in contrast to traditionally studied non-equilibrium systems in physics where the displacement from equilibrium is fixed [29, 30].

## RESULTS

### An ecological model of thermodynamically-constrained nutrient cycles

Our model describes a self-sustaining ecosystem in which *S* microbial species collectively recycle a set of environmental resources through sets of *R* thermodynamically-constrained redox transformations. Each species in the ecosystem corresponds to a different metabolic type, depending on the subset of these *R* transformations it can perform to maintain itself. Individuals of each species *α* extract an energy flux 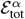 depending on the net energy released by coupling transformations, growing if they extract more than a prescribed maintenance energy *ε*_maint_, and dying if they extract less. Species dynamics modify the resource concentrations, making each transformation more or less thermodynamically favorable, and consequently changing the energy extracted by individuals of each species. Eventually, the ecosystem self-organizes to a steady state characterized by two sets of emergent quantities: (1) the abundances of each surviving species *N*_*α*_, where each individual extracts exactly *ε*_maint_, and (2) the fluxes of each of the *R* resource transformation cycles.

The key ingredient in our model — which distinguishes it from conventional “consumer-resource” models of ecosystems [26–28, 31–33] — is that resources are not single molecules with a certain energy content (Fig. 1a), but instead redox transformations (half-reactions) whose energy content is determined by electron and thermodynamic constraints (Fig. 1b). All *R* resources correspond to transformations *O*_*i*_ *↔R*_*i*_ between pairs of molecules *O*_*i*_, *R*_*i*_, representing the oxidized and reduced forms. A redox tower orders all resource pairs (*O*_*i*_, *R*_*i*_) by their chemical potential *μ*_*i*_, from least energetically favorable *O*_*i*_ *↔R*_*i*_ conversion (top) to most (bottom) (Fig. 1a). These chemical potentials *μ*_*i*_ are given by a standard state potential 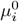 [8, 34] and an adjustment due to concentrations of, i.e., 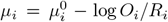. Each species *α* may exploit a specific subset of these transformations specified by kinetic coefficients *e*_*iα*_ ≥ 0; e.g., if *e*_*iα*_≠0, species *α* can transform *R*_*i*_ → *O*_*i*_ with kinetic coefficient *e*_*iα*_, releasing electrons at a potential *μ*_*i*_ which are then absorbed by another transformation *O*_*j*_ →*R*_*j*_ that the species participates in. The potential difference drop experienced by the electron is the energy available to this species. In practice, the electrons are transferred by an electron carrier (e.g., NADH) in the cell that is at potential *μ*_carrier,*α*_ intermediate to *μ*_*i*_ and *μ*_*j*_. We will assume there is only one electron carrier pool common to all redox transformations in species *α*.

**FIG. 1.**
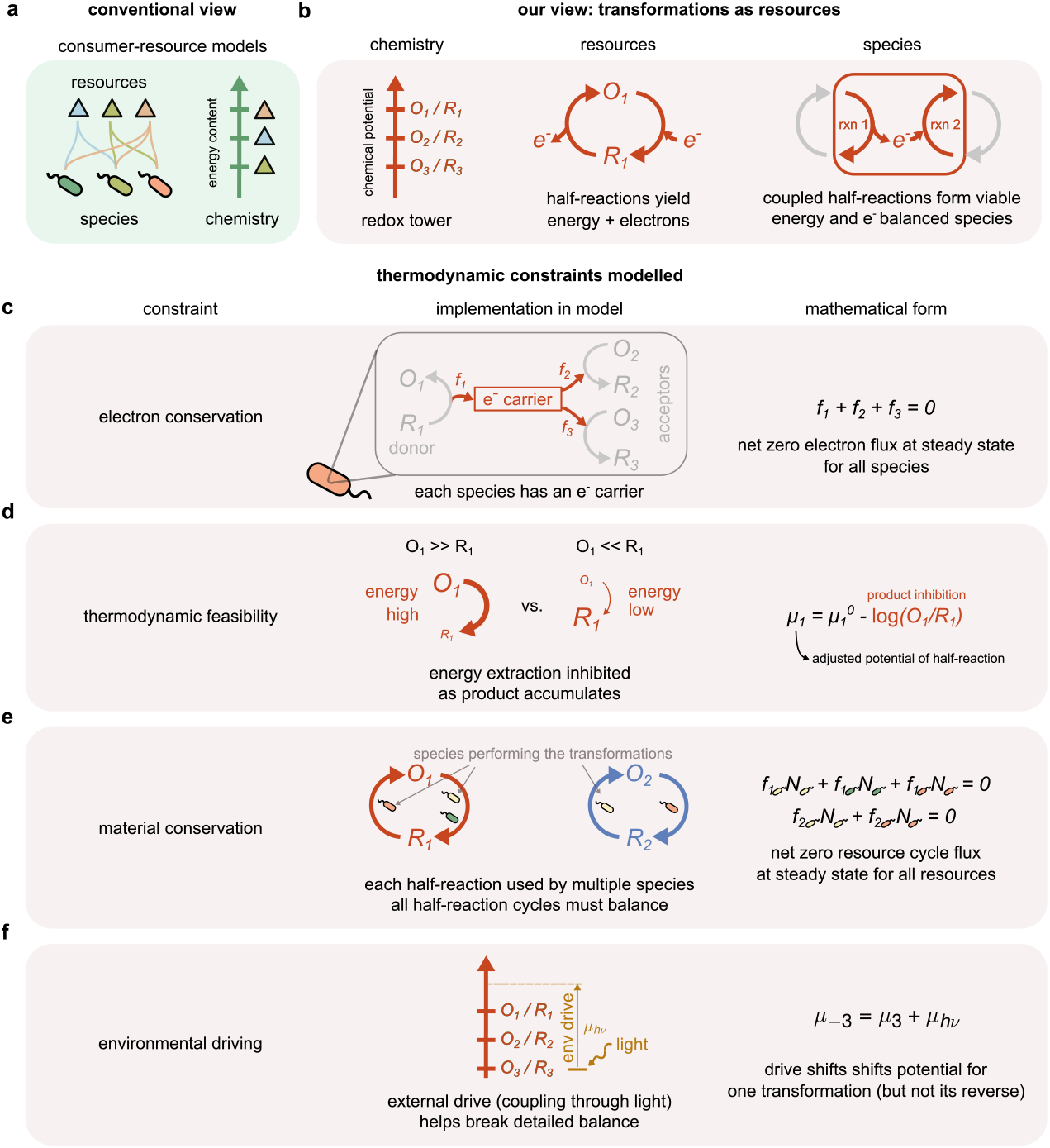
An ecological model of thermodynamically-constrained nutrient cycles. (a) In conventional models, species consume different molecules (triangles) that serve as resources with energy content. (b) In our model, resources are *transformations* of molecules, e.g., from an oxidized form *O*_*i*_ to their reduced form *R*_*i*_. Species gain energy by coupling transformations (or redox half-reactions) that donate electrons with transformations that accept electrons. Each (microbial) species is defined by the transformations it can carry out. Transformation fluxes must satisfy thermodynamic and conservation constraints in the ecosystem’s non-equilibrium steady state: (c) Electron conservation: The fluxes 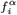 in different half-reactions carried out by a species *α* must sum to zero, i.e., 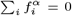 We assume that each microbial species has one electron carrier that shuttles electrons between the half-reactions. (d) Thermodynamic feasibility: The thermodynamic driving force (chemical potential) of each transformation, and consequently energy available, is calculated using thermodynamic principles, accounting for inhibition due to the concentrations of *O*_*i*_ and *R*_*i*_. (e) Material conservation: Fluxes in different parts of each molecular cycle (shown in red and blue), summed over contributions from all species, must be balanced. (f) Detailed balance breaking: Energy available by transforming matter in cycles is constrained by breaking of detailed balance. Detailed balance is broken by coupling some transformations (here, *O*_3_ → *R*_3_) to an energy source (e.g., light) which shifts their potential by *μ*_*hv*_ but does not affect the reverse transformation (*R*_3_ → *O*_3_).

Without any external energy source, the chemical potentials *μ*_*i*_ in the redox tower will obey detailed balance [29, 30]; i.e., as an electron is transferred in a loop through a cycle of redox transformations by different species, the net change in the electron’s energy must be zero. Consequently, in such a closed system, some species will have to *provide* energy to move the electron ‘uphill’ during metabolism, rendering such ecosystems not viable.

We will assume that detailed balance is broken across the redox tower because some transformations, say *R*_*j*_ →*O*_*j*_ are coupled to an external energy (but not matter) source (e.g., coupling the transformation H_2_O →O_2_ to sunlight during photosynthesis). Consequently, the chemical potential of *R*_*j*_ →*O*_*j*_ is shifted *μ*_*j*_ = *μ*_−*j*_ + *μ*_*hv*_ where *μ*_*j*_ is the chemical potential for the reverse transformation *O*_*j*_ →*R*_*j*_ (not coupled to light); the term *μ*_*hv*_ breaks detailed balance and is a key feature of the physical environment whose impact on ecosystems we will explore below.

The redox formalism allows us to correctly model the thermodynamic constraints on collective energy extraction. To maintain themselves at steady state, species in our model ecosystems must satisfy three constraints: electron conservation, material conservation and sufficient energy extraction. To conserve electrons, the total flux of electron donor and acceptor transformations must balance within each species (Fig. 1c, Methods). The per capita flux in transformation *O*_*i*_ →*R*_*i*_ due to species *α* is determined by species abundance *N*_*α*_, molecular concentration *O*_*i*_, and by thermodynamic driving forces Δ*μ*_*iα*_,

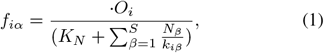

with *k*_*iα*_ determined by thermodynamic forces and fluxes (derivation in Supplementary Text),

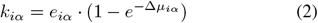

where Δ*μ*_*iα*_ is the change in potential of an electron released by transformation *O*_*i*_ → *R*_*i*_ and transferred to the electron carrier. Hence 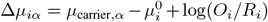 where 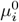 is the standard state chemical potential of the transformation *O*_*i*_ → *R*_*i*_ and *μ*_carrier,*α*_ is the chemical potential of the electron carrier for in species *α*.

These fluxes *f*_*iα*_ change the concentrations of *O*_*i*_, *R*_*i*_through the following dynamics:

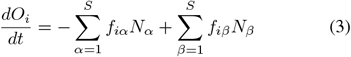

where *f*_*iα*_ is the flux of the transformation *O*_*i*_ →*R*_*i*_ performed by an individual of species *α* (as given in equation (1)), and *f*_*iβ*_ is the flux of the transformation *R*_*i*_→ *O*_*i*_ (similar to equation (1), but proportional to the reactant concentration *R*_*i*_, not *O*_*i*_ by individuals of species *β*. The first sum goes over all species transforming *O*_*i*_ →*R*_*i*_ and the second sum over species capable of the reverse. Similar equations hold for *R*_*i*_.

Each species *α* extracts a per capita energy flux 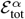 by coupling electrons between transformations at different potentials:

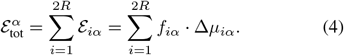

A species grows in abundance if this extracted energy exceeds a prescribed per capita maintenance energy *ε*_maint_:

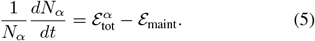

Finally, to conserve materials, as species in the ecosystem couple different half-reactions and transform resources from one form to another, all resource cycles must be balanced (Fig. 1e). Together, these constraints can be summarized as:

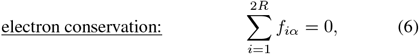

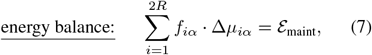

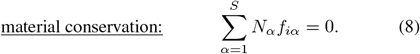

The last equation amounts to assuming that the ecosystem is fully closed to matter, and open only to an external source of energy (here, light energy *μ*_*hv*_) that breaks detailed balance for the chemical potentials *μ*_*i*_. While we use a fully materially closed ecosystem as an extreme case to illustrate our model (Figs. S1 and S2), many of our results hold for partially closed ecosystems as well, where some of the resources can be exchanged with the environment and equation (6) is modified (Fig. S3).

In addition to providing energy through transformations, matter also directly contributes to biomass [35]. We assume that the amount of matter sequestered as biomass is insignificant compared to the total availability (*O*_*i*_ + *R*_*i*_) and only focus on energy in this work. We leave an analysis of the dual role of matter in providing energy (through transformations) and biomass to future work.

The constraints encoded in our model naturally give rise to multiple solutions — there is a large space of ecosystems that satisfy them. To sample this space, we seeded a chosen physical environment with random mixtures of species with random metabolic strategies, and numerically evolved the dynamical equations until a subset of species found a stable composition or went extinct. Each ecosystem was provided with the same ‘physical environment’, i.e., concentrations of *R*_*i*_, *O*_*i*_ and energy input *μ*_*hv*_, but with a different initial set of *S* species chosen randomly and allowed to reach steady state (Methods). By simulating 1,000 such ecosystems, we sampled a large space of self-organized ecosystems.

### Emergent similarity of nutrient cycles

To quantify the size of this space of ecosystems, we measured their structure (set of species abundances) and function (set of resource fluxes *ϕ*_*i*_) and total energy extracted 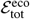) (Fig. 2c). Each point in structure space represents the abundances of each of the *S*_pool_ = 100 species in the pool in one of our simulated ecosystems (one species per dimension). Similarly, each point in flux space represents the fluxes in each of the *R* = 3 resource cycles in that ecosystem, and each point in the distribution of energy extracted represents the total energy 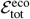captured from the external energy source by the ecosystem. To compare the size of the species and flux spaces, we projected them both onto a common two-dimensional space while preserving pairwise distances between ecosystems and computed the resulting area (see Methods); such a projection allows for a fair comparison of the convergence in species abundances and fluxes which are of different dimensionalities. We find that ecosystem-wide fluxes show less variability than the structure of species enabling these fluxes (Fig. 2d); further, while the variability in species abundances rapidly grows with the number of added species, the flux variability does not (Fig. 2e and Fig. S8). While alternative methods of comparing flux and species variability can change their absolute numbers, we expect this trend of increasing functional convergence with species diversity to be robust.

**FIG. 2.**
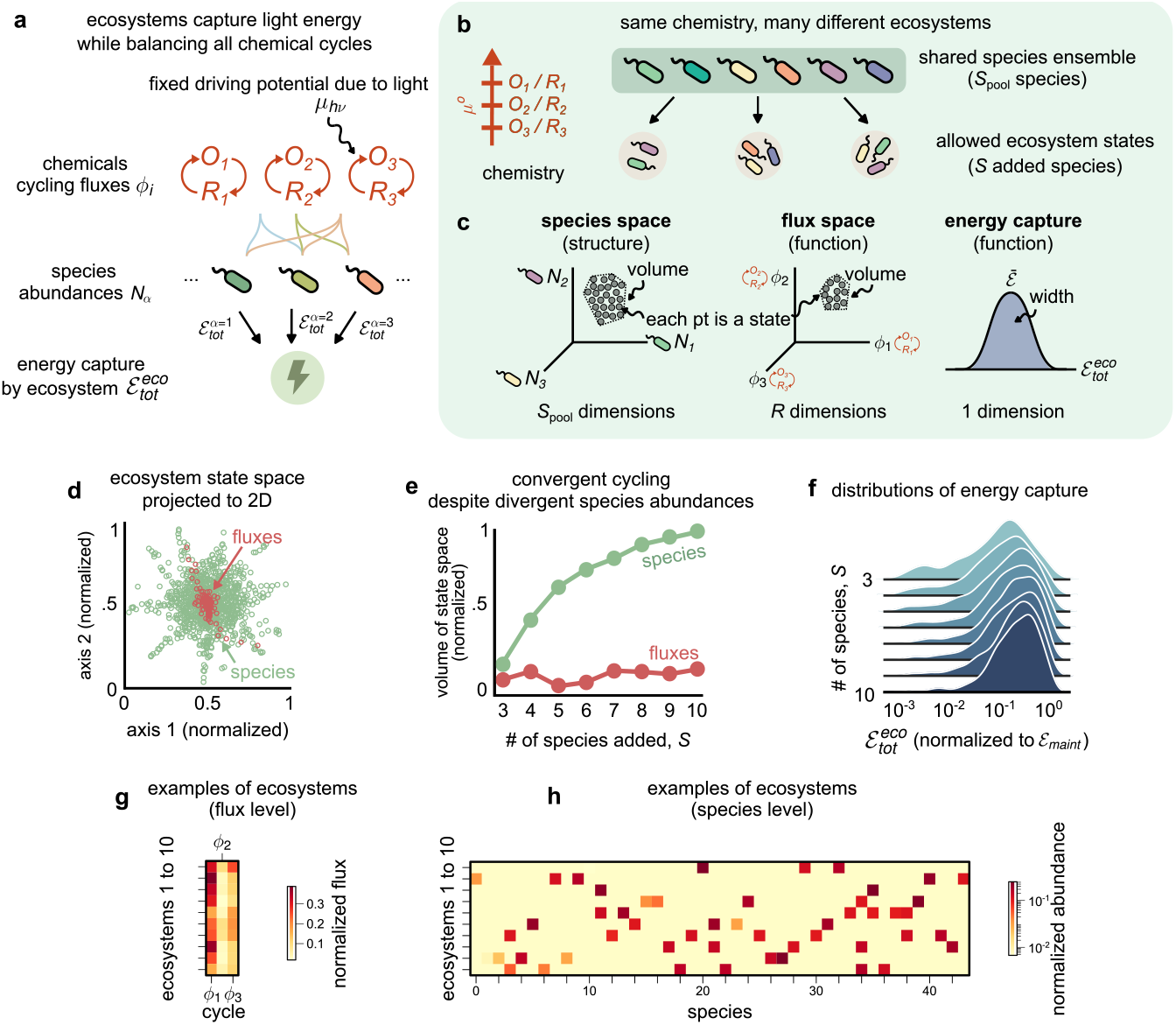
Ecosystems spontaneously implement correlated nutrient cycles and convergent energy extraction while containing distinct species. (a) Schematic showing how ecosystems in our model extract energy from externally supplied light (with driving potential *μ*_*hv*_), which affects the redox potentials of certain half-reactions. At steady state, all resources (*Oi, Ri*) are cycled with fluxes *fi*. Each microbial species (colored ellipses with wiggles) with abundances *Nα* carries out a subset of half-reactions (undirected colored links). By performing metabolic transformations, each surviving individual must extract a maintenance energy 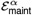 collectively, the entire ecosystem extracts an energy flux 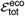 (b) Schematic showing a given physical environment consisting of *R* = 3 redox transformations (resources), and a pool of *S*_pool_ = 100 species (top rectangle) used to randomly assemble constraint-satisfying ecosystems in simulations. (c) Cartoon showing possible ecosystem solutions in a space of species abundances (left), resource cycle fluxes (middle), and a distribution of collective energy extracted (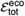 right). Scatter plot showing the space of species (green) and fluxes (red) from 1,000 randomly assembled ecosystems from simulation, projected to two dimensions using multi dimensional scaling (MDS) (Methods). (e) Line plot showing how the volume of the species (green) and flux (red) spaces scales with the number of species added, *S*, in assembled ecosystems. The flux space volume grows much slower than species space volume, indicating convergence in the function (fluxes) of constraint-satisfying ecosystems. (f) Distributions of the total energy extracted by ecosystems, 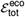, as a function of *S*. As ecosystems become more species-rich, ecosystems extract greater average energy with greater convergence (lower variance). (g–h) Heatmaps showing examples from 10 of the 1,000 randomly assembled ecosystems in (d–f), showing the (g) fluxes and (h) species abundances in detail. Each row shows an ecosystem, while each column shows a resource (in (g)) and a species (in (h)).

A nearly constant flux variability, despite increasing species variability, suggests that fluxes are a convergent feature of ecosystems that perform thermodynamically-constrained transformations. Indeed, the convergence of fluxes despite species divergence is evident from a few examples of our model ecosystems (Fig. 2g–h, fluxes and species respectively). We also found that ecosystems had more convergent structure when coarse-grained by metabolic type (the set of transformations each species could perform) than at the species level (Fig. S4), similar to what has been observed in metagenomic data from field surveys of microbial communities [31, 36, 37].

While functional convergence of fluxes and metabolic types has been noted in other contexts, both in experiments[36, 38] and from other models[39, 40], our model allows us to go further and study convergence of explicitly thermodynamic functions of ecosystems. We find that the distributions of total energy extracted by our model ecosystems also converges, with the mean energy extracted increasing and variance in energy extracted decreasing with diversity, before ultimately saturating (Fig. 2f). Finally, we determined the energy contribution *ε*_*ij*_ of each pair (*i, j*) of redox transformations to the total energy harvested; we found that such global ‘redox strategies’ of ecosystems became progressively similar with increasing diversity (Fig. S5).

Thus, we find that thermodynamic constraints of coupled transformations self-organize ecosystems such that collective functions – here, the distribution of fluxes and energy extracted across cycles – are similar across distinct ecosystems that differ widely in their structure (species content).

### Collective functions become more variable with stronger environmental driving

The influence of the physical environment (e.g., nutrients supplied) is at the core of all ecology; but the impact of thermodynamic properties of the environment on ecosystem organization has been studied only in models of specific systems (e.g., communities with hydrogenotrophic methanogens where end-product inhibition is crucial [41–43]).

In our model, the physical environment consists of a redox tower with standard-state chemical potentials 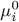 of the redox transformations *R*_*i*_ *↔ O*_*i*_ and the total amounts of matter *R*_*i*_ + *O*_*i*_ available for each transformation. Most critically, the environment also includes an external energy drive such as light that shifts the potential of one redox transformation by *μ*_*hv*_ (but does not affect the reverse transformation).

Without such an external drive *μ*_*hv*_, transformations within an ecosystem are constrained to satisfy detailed balance, a defining property of equilibrium systems, and thus no net energy can be extracted [29, 30]. At the same time, non-equilibrium driving by *μ*_*hv*_ does not by itself set the energy extracted by the ecosystem; species abundances must self-organize and ultimately determine the energy extracted. Fixing the environmental driving potential *μ*_*hv*_ is distinct from fixing the flux of an external resource which has been the focus of previous modeling approaches [27, 39, 44]. In an electrical circuit analogy, *μ*_*hv*_ sets the external voltage while prior approaches typically set an external current. These two approaches are especially distinct in self-organized ecosystems where the analog of ‘resistance’ is not fixed since species abundances evolve through population dynamics.

We sought to understand how external drive *μ*_*hv*_ in the physical environment affects the number of viable ecosystems, and their functional convergence. We simulated an ensemble of ecosystems for each of several environmental driving potentials *μ*_*hv*_. We found that ecosystems can sustain themselves only if *μ*_*hv*_ exceeds the smallest potential difference Δ*μ*_min_ (Fig. 3b) between all potentials *μ*_*i*_ on the redox tower (Fig. 3a). Below this level, ecosystems were energy deprived due to insufficient environmental driving (Fig. 3b, gray region) and not viable.

**FIG. 3.**
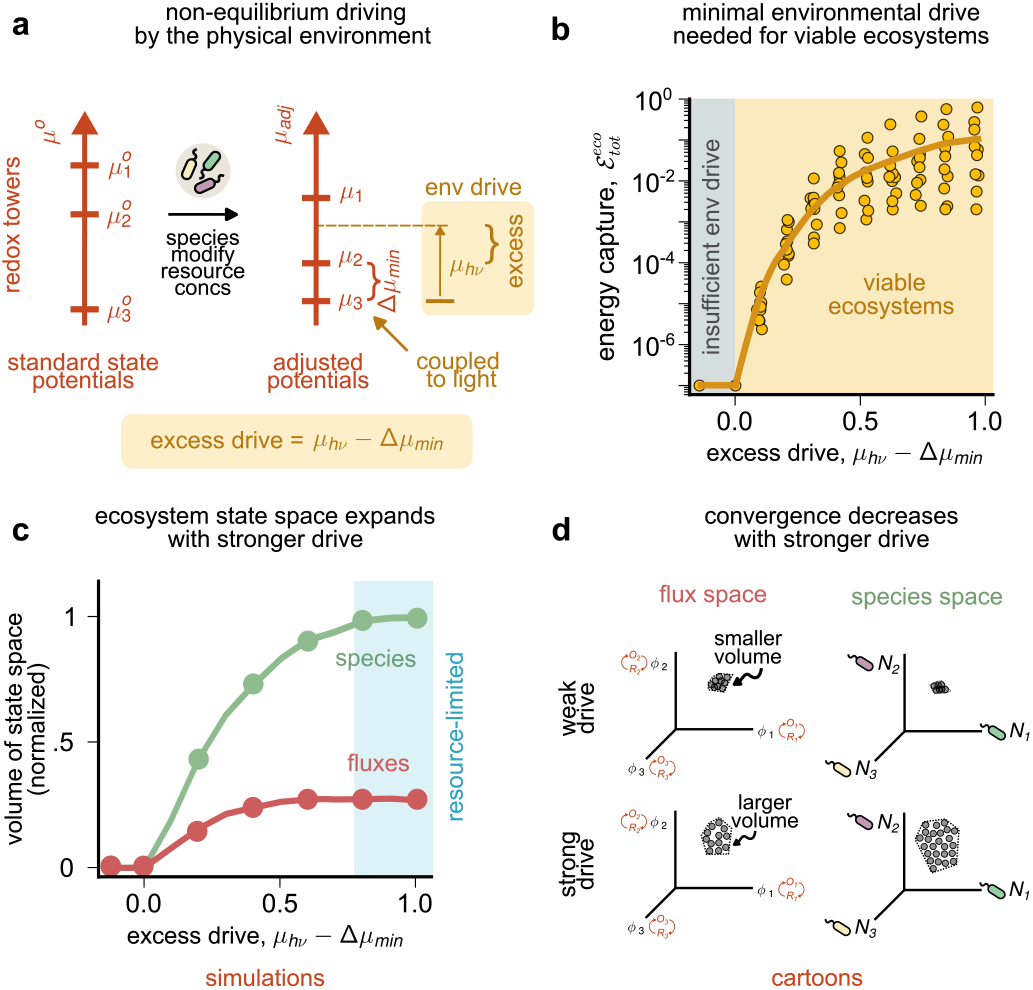
Nutrient cycles and energy extraction become more variable with stronger environmental driving. (a) Schematic showing non-equilibrium driving by the environment, modeled as the coupling of one of the half-transformations (at the bottom of the redox tower) to light energy. Environmental driving shifts the chemical potential of of the transformation *R*_3_ → *O*_3_ by *μ*_*hv*_, but not the reverse, thus breaking detailed balance. For ecosystems to be viable and satisfy all the modeled thermodynamic constraints, the environmental drive *μ*_*hv*_ must exceed the smallest adjusted potential difference Δ*μ*_min_ between *O*_3_*/R*_3_ and the closest redox pair. This minimal drive requirement is a statement about adjusted potentials on the redox tower, not the standard state potentials. The former depend on the species present and their abundances, and are thus self-organized. We quantify *μ*_*hv*_− Δ*μ*_min_ as the extent of environmental drive beyond the minimal (Δ*μ*_min_) needed for a viable ecosystem. (b) Scatter plot showing the energy extracted by ecosystems from our model as a function of excess drive. Each point represents a random self-organized ecosystem simulated using our model, under the same conditions as in Fig. 2, at varying *μ*_*hv*_. The solid line shows a moving average. No ecosystems are viable without sufficient environmental driving (gray region). Increasing *μ*_*hv*_ increases energy extraction on average (mustard region). (c) Line plot showing the volume of ecosystem state space in species abundances (green) and resource fluxes (red), as in Fig. 2e, but with varying excess drive *μ*_*hv*_ − Δ*μ*_min_. Ecosystems near equilibrium are strongly constrained; the volume of viable ecosystems grows rapidly with stronger environmental driving, eventually saturating at large *μ*_*hv*_. Results are shown from simulations with 1,000 randomly assembled ecosystems. (d) Cartoons showing how flux (red) and species (green) spaces expand (decreasing convergence) with stronger drive, with spaces represented as in Fig. 2c.

Note that the minimal environmental drive Δ*μ*_min_ for ecosystem viability depends on redox potentials *μ*_*i*_ that account for product inhibition in the ecosystem self-organized by the species present and not on standard state potentials 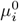 Thus, in distinction to prior work, our model can determine the minimal environmental drive for ecosystem viability as a function of which species are present (Fig. S6).

For strong enough driving *μ*_*hv*_ *>* Δ*μ*_min_, ecosystems self-organized to extract greater collective energy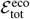 with increasing external chemical potential *μ*_*hv*_ (Fig. 2b, mustard region). The mean 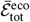 eventually saturated at large *hν*, due to limitation from total resource concentrations *Σi*(*O*_*i*_ + *R*_*i*_) (Fig. S7).

Notably, the variance in 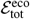also increased with *μ*_*hv*_ (Fig. 3b, yellow circles), suggesting that stronger environmental driving decreased convergence in the collective energy extracted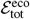. In addition, both the species and flux spaces of constraint-satisfying ecosystems decreased in size with decreased environmental driving (Fig. 3c, species volume in green, flux volume in red). The species and flux spaces, as well as collective energy extracted, are emergent features, representing how ecosystems self-organize once the physical environment drives them sufficiently far from equilibrium.

Together, these results suggest a relationship between non-equilibrium environmental driving and the degree of convergence in ecosystems. When ecosystems operate in environments that are closer to equilibrium (small *μ*_*hv*_), there is a smaller space of distinct ways to cycle all resources successfully and provide maintenance energy for all organisms. Consequently, convergence is strongest in environments that are near equilibrium.

### Near-optimal energy extraction by self-organized ecosystems

The collective energy extracted 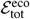, an emergent community function, is not explicitly optimized by any aspect of our model; our population dynamics equations model ‘greedy’ or ‘selfish’ biological species that grow in abundance to extract the most energy that each species can extract in the context of a self-sustaining ecosystem.

A natural question is how this energy extracted by an ecosystem of locally selfish replicators compares to an alternative community of ‘metabolic machines’ agents whose abundances are determined by global energy optimization but that are still subject to the same thermodynamic and flux constraints. We explored one such alternative framework based on communities of non-living ‘metabolic machines’ identical to self-replicating living species in terms of metabolism but not subject to birth-death population dynamics. More explicitly, each machine species *α* was identical to living species *α* in terms of its metabolic properties (e.g., 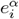 which dictate which redox transformations *i* species *α* can perform). Further, energy extraction by the machines were subject to all the thermodynamic, electron and material conservation constraints discussed earlier.

We initialized a community of machines with initial random abundances and adjusted those abundances minimally to obey electron and mass conservation constraints (see Methods). We computed the energy extracted by these communities of effectively random abundances (Fig. 4b, gray). We then evolved these random initial set of abundances in two distinct ways: (1) Global community-wide energy optimization for machines, (2) Local maintenance-energy based population dynamics for self-replicators.

**FIG. 4.**
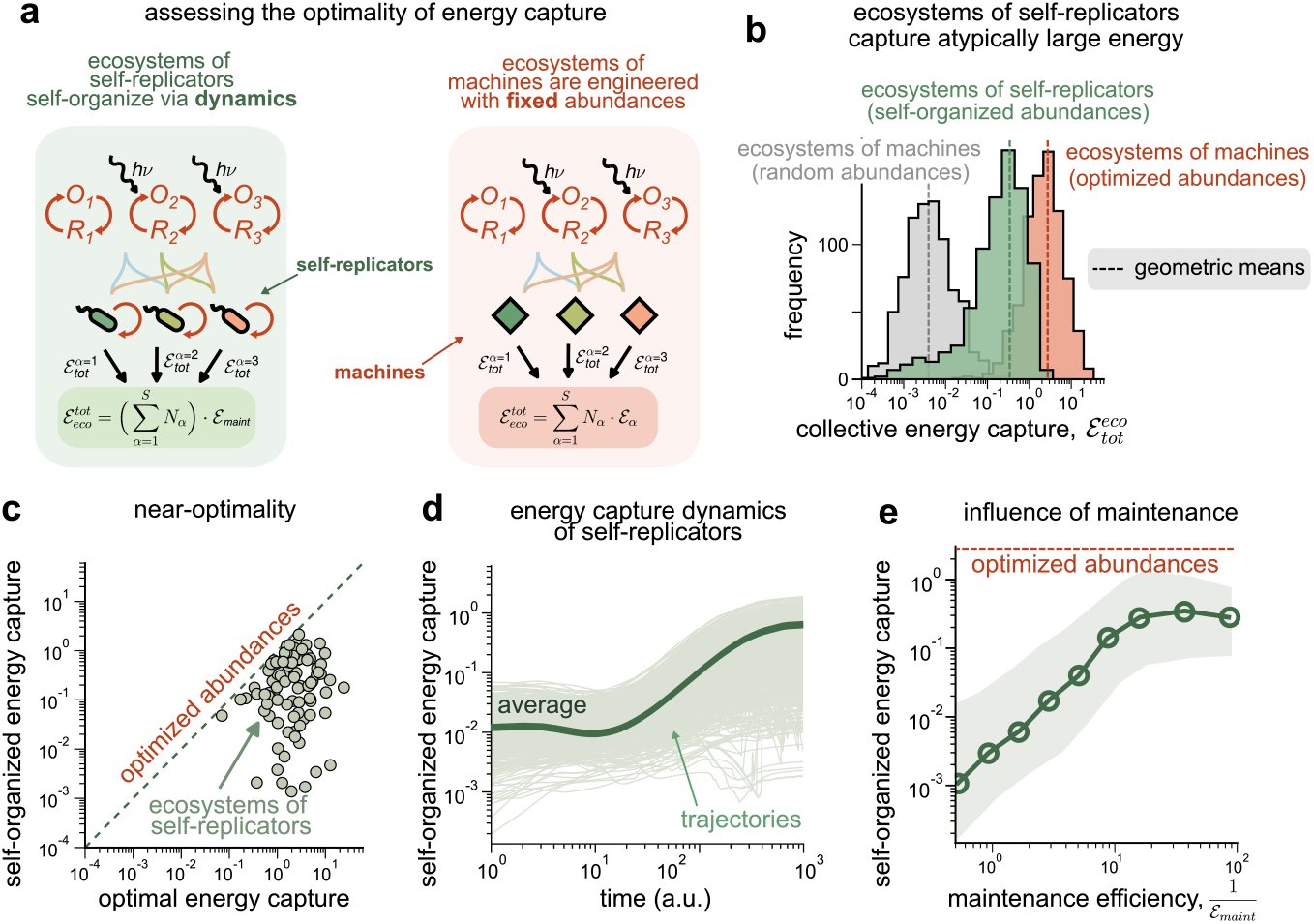
Ecosystems self-organize to extract a near optimal amount of energy. (a) We compare energy extracted by ecosystems of living organisms (self-replicators) to a ecosystem of machines with identical metabolic capabilities. Like living organisms, machines come in multiple species with distinct metabolic types and are subject to the same thermodynamic and conservation constraints. However, machines have fixed abundances, not subject to birth-death dynamics driven by maintenance energy requirements found in living systems. (b) Histograms of the total energy extracted 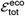by ecosystems of machines with random abundances (gray; representing initial conditions of ecosystem assembly), ecosystems of self-replicators with abundances self-organized by birth-death dynamics based on maintenance energy (green), and machines with abundances chosen to maximize 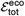 (orange). Ecosystems of self-replicators reach steady states that extract *∼* 100*×* more energy than initial random abundances. (c) Scatter plot showing the energy extracted 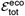 by ecosystems with self-organized abundances (*y*-axis) and optimized abundances (*x*-axis); both systems are constituted from the same pool of metabolic strategies. (d) Trajectories showing the dynamics of ecosystems of self-replicators reaching steady states over time (1,000 simulations). (e) Line plot showing how the average 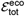 (solid green) changes as a function of the maintenance efficiency per cell 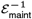 Each point represents the average collective energy extracted by ecosystems of self-replicators at the specified *ε*_maint_ simulated using our model (averaged over 1,000 simulations); the green envelope their s.e.m.; the red dashed line shows the geometric mean of the energy extracted by ecosystems of machines with optimized abundances.

1. Global optimization for machines: we optimized the abundances *N*_*α*_ of each species *α*, subject to electron and flux constraints to maximize community-wide energy extraction; the machines were not subject to any population dynamics (Methods and Fig. 4a). Consequently, a machine species of type *α* could exist at any abundance, dictated only by what was optimal for the community as a whole, even if its own energy extracted would have lead to higher or lower abundance (or even extinction) according to ‘selfish’ population dynamics (equation (6)). The energy, as expected, is dramatically higher after such global optimization (Fig. 4b, orange).
2. Local population dynamics for self-replicators: Starting from random initial abundances, we also ran population dynamics based on maintenance energy (equation (7)). Unlike global optimization, now each species grows in abundance ‘greedily’ until it cannot extract an energy that exceeds maintenance energy. Despite such local greedy evolution, we found that on average, the mean of the energy distribution for self-replicating agents was 100*×*higher than the mean for random abundances and only 10*×* lower than the mean for globally optimized community of machines (Fig. 4b). This result suggests that ‘selfish’ population dynamics drives random initial abundances most of the way to abundances predicted by a global optimization algorithm, even though the ‘selfish’ dynamics are only aware of the energy extracted by each species and do not explicitly try to maximize global energy extraction. Atypically large energy extraction by ecosystems was true for nearly every set of self-replicating (living) species tested (Fig. 4c), suggesting that the difference between means was not driven by only a few extremely efficient ecosystems. The population dynamics of each species naturally drove ecosystems towards such atypically large energy extraction, with 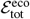 on average increasing over the dynamical trajectories of ecosystems (Fig. 4d).

Finally, since the population dynamics of living cells (but not machines) were affected by maintenance energy *ε*_maint_, we studied how the collective energy extracted depends on it. Our simulations revealed that the total energy extracted is relatively independent of *ε*_maint_ for low *ε*_maint_ but falls at high*ε* _maint_. At high *ε*_maint_, species abundances *N*^*α*^ are the limiting factor for fluxes in the redox transformations. But as *ε*_maint_ is reduced, species abundances increase and eventually, fluxes are limited by the amount of resources *R*_*i*_ + *O*_*i*_ and not by species abundances; consequently, the collective energy extracted 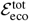 is relatively constant.

For ecosystems of machines, collective energy extracted depends on both the energy extracted by each individual machine, as well as the number of individuals 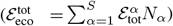 unlike in ecosystems of self-replicators where it depends only on total biomass 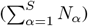 since each individual is constrained to extract maintenance energy 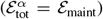 at steady state. Hence, machine ecosystems may arrange species abundances to extract more energy collectively, even though the distribution of per capita energy extracted across species may be quite broad.

Taken together, ecosystems of self-replicators with low maintanence energy *ε*_maint_ capture energy comparable to ecosystems of machines with globally optimized abundances. The difference in energy extracted between self-organized and globally optimized ecosystems quantify a “greedy gap” of 10*×* on average, i.e., the extent to which greedy self-replication constrains collective energy extracted in ecosystems of living cells.

## DISCUSSION

Here, we proposed a new theoretical framework to study self-organized energy extraction by ecosystems, which incorporates an essential aspect of metabolism thus far missing from most ecological models: organisms acquire energy through redox transformations of matter, not matter itself. Modeling resources as transformations is not only biologically accurate but also provides a modeling framework to address fundamental questions about thermodynamic constraints on the self-organization of ecosystems.

Using this model, we studied the impact of closure to matter and redox metabolism on nutrient cycling in ecosystems. By sampling large random ensembles that satisfy these constraints, we found that ecosystems converge in similar thermodynamic features, such as nutrient cycling fluxes and the total energy extracted. We also found that ecosystems converge to a lesser degree when the extent of detailed balance breaking increases, i.e., when the available potential energy from light increases. We also find that the collective energy extraction is remarkably high, given that dynamics through which ecosystems assemble in our model involve local ‘selfish’ growth rules with no awareness of the global collective energy extracted. That ecosystems extracted atypically large energy — 100-fold closer to the theoretical maximum when compared with ecosystems with random abundances — from their external environment (light) is an emergent feature of the dynamics of our model, arising purely from basic constraints introduced by acknowledging redox transformations and closure to matter in ecosystems.

While there is a growing body of work documenting functional convergence in ecosystems [31, 36–3 our work expands the domain of functional convergence to new, explicitly thermodynamic features and the impact of the environmental driving potential. Our basic result is that functional convergence is strongest for weakly-driven near-equilibrium ecosystems; strong external driving potentials decrease convergence. This result suggests a deep theoretical connection between different manifestations of functional convergence as representing the multiplicity of ways in which communities break detailed balance.

To calculate the energy extracted, we made the simplifying assumption that organisms extracted energy equal to the entire energy gap between their respective donors and acceptors. Realistically, a fraction of the energy gap is extracted and stored as chemical energy (e.g., ATP) while the rest is dissipated [34, 45]. This can be incorporated in our frame-work, by including suitable “ATP coupling” parameters for every donor-acceptor pair. We expect that such an extension of our model doing will generally increase niche competition between species coupling the same donor-acceptor pairs, and thus might decrease total energy extraction.

While this manuscript focused on materially closed, energy-limited ecosystems, our theoretical framework can be extended in a variety of ways. Examples include extending the framework to account for the dual role of resources, as sources of both energy and biomass, where organismal growth would depend on which of the two — energy extraction or biomass generation — are limiting. Another is to extend the model to be spatially explicit, in order to study spatio-temporal pattern formation such as self-organized stratification in microbial mats and Winogradsky columns. Finally, our work can help identify likely signatures of life on redox towers in astrobiological contexts, e.g., by studying the self-organized adjusted potentials (like in Fig. 3a and Fig. S9).In all these questions, redox constraints are essential to the underlying phenomena. Thus our work opens up new lines of inquiry in redox ecology.

In addition to ecology, our framework could be used to understand the role of energy extracted in driving the evolution of living systems. Ecologists have widely argued that energy might serve as a natural fitness function during the evolution of biological communities [46]. However, natural selection acts on individuals and not on directly on community function. Extensions of our work can provide a framework to understand the tension between ‘selfish’ evolution of individuals and collective energy extracted by an ecosystem. Such a framework could be useful in guiding the engineering of evolutionarily stable photosynthetic communities.

Another feature of our model is *emergent* detailed balance breaking. Unlike in other models of non-equilibrium systems [29]— where the extent of detailed balance breaking is a fixed external quantity — in our work, we set a fixed external driving potential (e.g., that of light) in the redox tower but the amount of detailed balance breaking is determined by self-organization of the ecosystem (e.g., through species abundances and material abundances that change chemical potentials through product inhibition). As a consequence, e.g., there is a minimal non-zero external drive below which there is no detailed balance breaking (Fig. 3b, gray region). In this way, our work suggests an ecology-inspired framework for studying the emergence of spontaneously self-organized non-equilibrium steady states (NESS), adding to prior work on the origin of dissipative structures inspired by Rayleigh-Benard convection cells and other physical systems [47–49].

## METHODS

Please see SI Appendix for detailed materials and methods. Briefly, we constructed a theoretical framework for microbial ecosystems, whose constituent species extract energy through thermodynamically-constrained redox conversions of matter. Our theory relied on fundamental physico-chemical constraints which we outlined in Results and further elaborate on in the SI Appendix. We then derived a dynamical model of ecosystem self-organization based on the constraints outlined in equations (4)-(6), resulting in equations (7) and (8) in the main text by using thermodynamically accurate expressions for process rates (e.g., product inhibition and energy-dependent forcing). Using numerical simulations of these dynamical equations, we generated ensembles of ecosystems at steady state with a fixed physical environment, but different initial sets of species (Fig. 2), and with varying levels of detailed balance breaking *μ*_*hv*_ (Fig. 3). We then used numerical optimization and root-finding techniques to generate analogous ecosystems of non-living machines, by finding solutions of the constraint equations (4) and (6) which globally maximized the collective energy extracted given by equation (3) (Fig. 4).

## ACKNOWLEDGEMENTS

We thank the Kavli Institute for Theoretical Physics, UCSB, where this project took root. We thank G. Birzu, O.X. Cordero, W. W. Fischer, J. Goldford, S. Kuehn, S. Maslov, A. Narla, and M. Tikhonov for discussions. This research was supported in part by the National Science Foundation under grant number NSF PHY-1748958. A.G. is supported by the Gordon and Betty Moore Foundation as a Physics of Living Systems Fellow through grant number GBMF4513. crobiology **75**, 501 (2006).

## References

[1] E. P. Odum, Fundamentals of ecology, 3rd, Edition, Philadephia: WB Saunders (1971).

[2] C. J. Cleveland, R. Kaufmann, and T. Lawrence, Fundamental principles of energy, Encyclopedia of Earth, 79 (2008).

[3] M. L. Messager, B. Lehner, G. Grill, I. Nedeva, and O. Schmitt, Estimating the volume and age of water stored in global lakes using a geo-statistical approach, Nature communications 7, 13603 (2016).

[4] J. P. Grotzinger and A. H. Knoll, Stromatolites in precambrian carbonates: evolutionary mileposts or environmental dipsticks?, Annual review of earth and planetary sciences 27, 313 (1999).

[5] G. A. Zavarzin, Winogradsky and modern microbiology, Microbiology 75, 501 (2006).

[6] E. Pagaling, K. Vassileva, C. G. Mills, T. Bush, R. A. Blythe, J. Schwarz-Linek, F. Strathdee, R. J. Allen, and A. Free, Assembly of microbial communities in replicate nutrient-cycling model ecosystems follows divergent trajectories, leading to alternate stable states, Environmental microbiology 19, 3374 (2017).

[7] L. M. de Jesùs Astacio, K. H. Prabhakara, Z. Li, H. Mickalide, and S. Kuehn, Closed microbial communities self-organize to persistently cycle carbon, Proceedings of the National Academy of Sciences 118, e2013564118 (2021).

[8] T. D. Brock, M. T. Madigan, J. M. Martinko, and J. Parker, Brock biology of microorganisms (Upper Saddle River (NJ): Prentice-Hall, 2003., 2003).

[9] P. G. Falkowski, T. Fenchel, and E. F. Delong, The microbial engines that drive earth’s biogeochemical cycles, science 320, 1034 (2008).

[10] B. B. Jørgensen and D. J. Des Marais, Competition for sulfide among colorless and purple sulfur bacteria in cyanobacterial mats, FEMS Microbiology Ecology 2, 179 (1986).

[11] D. E. Canfield and D. J. Des Marais, Biogeochemical cycles of carbon, sulfur, and free oxygen in a microbial mat, Geochimica et Cosmochimica acta 57, 3971 (1993).

[12] M. C. Rillig and J. Antonovics, Microbial biospherics: The experimental study of ecosystem function and evolution, Proceedings of the National Academy of Sciences 116, 11093 (2019).

[13] D. J. Esteban, B. Hysa, and C. Bartow-McKenney, Temporal and spatial distribution of the microbial community of winogradsky columns, PLoS One 10, e0134588 (2015).

[14] T. Bush, I. Butler, A. Free, and R. Allen, Redox regime shifts in microbially mediated biogeochemical cycles, Biogeosciences 12, 3713 (2015).

[15] J. A. Christie-Oleza, D. Sousoni, M. Lloyd, J. Armengaud, and D. J. Scanlan, Nutrient recycling facilitates long-term stability of marine microbial phototroph–heterotroph interactions, Nature microbiology 2, 1 (2017).

[16] F. B. Taub and A. K. McLaskey, Pressure, o2, and co2, in aquatic closed ecological systems, Advances in Space Research 51, 812 (2013).

[17] K. Matsui, S. Kono, A. Saeki, N. Ishii, M.-G. Min, and Z. i. Kawabata, Direct and indirect interactions for coexistence in a species-defined microcosm, Hydrobiologia 435, 109 (2000).

[18] F. B. Taub, Closed ecological systems, Annual Review of Ecology and Systematics 5, 139 (1974).

[19] D. Obenhuber and C. Folsome, Carbon recycling in materially closed ecological life support systems, Biosystems 21, 165 (1988).

[20] F. Kracke, I. Vassilev, and J. O. Krömer, Microbial electron transport and energy conservation–the foundation for optimizing bioelectrochemical systems, Frontiers in microbiology 6, 575 (2015).

[21] U. F. Lingappa, N. T. Stein, K. S. Metcalfe, T. M. Present, V. J. Orphan, J. P. Grotzinger, A. H. Knoll, E. J. Trower, M. L. Gomes, and W. W. Fischer, Early impacts of climate change on a coastal marine microbial mat ecosystem, Science Advances 8, eabm7826 (2022).

[22] T. W. Lyons, C. T. Reinhard, and N. J. Planavsky, The rise of oxygen in earth’s early ocean and atmosphere, Nature 506, 307 (2014).

[23] J. Imbrie and K. P. Imbrie, Ice ages: solving the mystery (Harvard University Press, 1986).

[24] P. F. Hoffman, A. J. Kaufman, G. P. Halverson, and D. P. Schrag, A neoproterozoic snowball earth, science 281, 1342 (1998).

[25] J. F. Kasting, Bolide impacts and the oxidation state of carbon in the earth’s early atmosphere, Origins of Life and Evolution of the Biosphere 20, 199 (1990).

[26] A. Goyal and S. Maslov, Diversity, stability, and reproducibility in stochastically assembled microbial ecosystems, Physical Review Letters 120, 158102 (2018).

[27] R. Marsland III, W. Cui, J. Goldford, A. Sanchez, K. Korolev, and P. Mehta, Available energy fluxes drive a transition in the diversity, stability, and functional structure of microbial communities, PLoS computational biology 15, e1006793 (2019).

[28] R. MacArthur, Species packing and competitive equilibrium for many species, Theoretical population biology 1, 1 (1970).

[29] H. Qian, Phosphorylation energy hypothesis: open chemical systems and their biological functions, Annu. Rev. Phys. Chem. 58, 113 (2007).

[30] T. L. Hill, Free energy transduction and biochemical cycle kinetics (Courier Corporation, 2013).

[31] J. E. Goldford, N. Lu, D. Bajić, S. Estrela, M. Tikhonov, A. Sanchez-Gorostiaga, D. Segre, P. Mehta, and A. Sanchez, Emergent simplicity in microbial community assembly, Science 361, 469 (2018).

[32] M. Tikhonov and R. Monasson, Collective phase in resource competition in a highly diverse ecosystem, Physical review letters 118, 048103 (2017).

[33] A. Posfai, T. Taillefumier, and N. S. Wingreen, Metabolic tradeoffs promote diversity in a model ecosystem, Physical review letters 118, 028103 (2017).

[34] Q. Jin and C. M. Bethke, A new rate law describing microbial respiration, Applied and Environmental Microbiology 69, 2340 (2003).

[35] M. Scott, C. W. Gunderson, E. M. Mateescu, Z. Zhang, and T. Hwa, Interdependence of cell growth and gene expression: origins and consequences, Science 330, 1099 (2010).

[36] S. Louca, S. M. Jacques, A. P. Pires, J. S. Leal, D. S. Srivastava, L. W. Parfrey, V. F. Farjalla, and M. Doebeli, High taxonomic variability despite stable functional structure across microbial communities, Nature ecology & evolution 1, 0015 (2016).

[37] S. Louca, M. F. Polz, F. Mazel, M. B. Albright, J. A. Huber, M. I. O’Connor, M. Ackermann, A. S. Hahn, D. S. Srivastava, S. A. Crowe, et al., Function and functional redundancy in microbial systems, Nature ecology & evolution 2, 936 (2018).

[38] S. Estrela, J. C. Vila, N. Lu, D. Bajić, M. Rebolleda-Gómez, C.-Y. Chang, J. E. Goldford, A. Sanchez-Gorostiaga, and A. Sánchez, Functional attractors in microbial community assembly, Cell Systems 13, 29 (2022).

[39] A. B. George, T. Wang, and S. Maslov, Functional universality in slow-growing microbial communities arises from thermodynamic constraints, arXiv preprint 2203.06128 (2022).

[40] L. Fant, I. Macocco, and J. Grilli, Eco-evolutionary dynamics lead to functionally robust and redundant communities, bioRxiv, 2021 (2021).

[41] T. Lynch, Y. Wang, B. van Brunt, D. Pacheco, and P. Janssen, Modelling thermodynamic feedback on the metabolism of hydrogenotrophic methanogens, Journal of Theoretical Biology 477, 14 (2019).

[42] H. Delattre, J. Chen, M. J. Wade, and O. S. Soyer, Thermodynamic modelling of synthetic communities predicts minimum free energy requirements for sulfate reduction and methanogenesis, Journal of the Royal Society Interface 17, 20200053 (2020).

[43] C.-Y. Hoh and R. Cord-Ruwisch, A practical kinetic model that considers endproduct inhibition in anaerobic digestion processes by including the equilibrium constant, Biotechnology and bioengineering 51, 597 (1996).

[44] J. Cook, S. Pawar, and R. G. Endres, Thermodynamic constraints on the assembly and diversity of microbial ecosystems are different near to and far from equilibrium, PLOS Computational Biology 17, e1009643 (2021).

[45] Q. Jin and C. M. Bethke, Predicting the rate of microbial respiration in geochemical environments, Geochimica et Cosmochimica Acta 69, 1133 (2005).

[46] A. J. Lotka, Contribution to the energetics of evolution, Proceedings of the National Academy of Sciences 8, 147 (1922).

[47] D. Kondepudi and I. Prigogine, Modern thermodynamics: from heat engines to dissipative structures (John Wiley & Sons, 2014).

[48] J. L. England, Statistical physics of self-replication, The Journal of chemical physics 139, 09B623 1 (2013).

[49] J. L. England, Dissipative adaptation in driven self-assembly, Nature nanotechnology 10, 919 (2015).

