## Supplementary Materials for "Closed ecosystems extract energy through self-organized nutrient cycles"


Akshit Goyal

*Department of Physics, Massachusetts Insitute of Technology, Cambridge, MA 02139.*

Avi I. Flamholz

*Division of Biology and Biological Engineering,*

*California Institute of Technology, Pasadena, CA 91125.*

Alexander P. Petroff

*Department of Physics, Clark University, Worcester, MA 01610.*

Arvind Murugan

*Department of Physics, University of Chicago, Chicago, IL 60637.*


### I. DESCRIPTION OF OUR MATHEMATICAL MODEL

As briefly described in the main text, our model describes a dynamical closed ecosystem in which $S$ microbial species collectively recycle a set of environmental resources through sets of $R$ thermodynamically-constrained redox transformations. Each species in the ecosystem corresponds to a different metabolic type, depending on the subset of these $R$ transformations it can perform to maintain itself. We track the dynamics of species abundances, $N_\alpha$ and resource molecule concentrations $R_i$, and study the properties of steady states of this system. For simplicity, we assume fast transport kinetics across microbial species, so that the intracellular concentrations of molecules that each species can transform is the same as the extracellular concentration. This allows us to only track the dynamics of only one set of concentrations for each molecule. Further, we assume that species dynamics are driven only by energy requirements, without any biomass turnover. This assumption allows us to study energy extraction at steady state, which for each surviving species must equal a prescribed maintenance energy $\mathcal{E}_{maint}$ (a tunable parameter of our model, studied in Fig. 4e). Thus, growing microbial species are limited only by energy, and we ignore the requirement of biomass building blocks. Similarly, we ignore the accumulation of organic matter by dying species in this work, and aim to study that model in greater detail in later work. We will now describe the model in greater detail, reproducing some sections from the main text description for the sake of completeness.

Each microbial species extracts energy through redox metabolic transformations (half-reactions) whose energy content is determined by electron and thermodynamic constraints (Fig. 1b). All $R$ resources correspond to a pair of transformations $O_i \leftrightarrow R_i$ between pairs of molecules $O_i, R_i$, representing the oxidized and reduced forms, respectively. A redox tower orders all resource pairs $(O_i, R_i)$ by their chemical potential $\mu_i$, from least energetically favorable $O_i \to R_i$ conversion (top) to most (bottom) (Fig. 1a).

These chemical potentials $\mu_i$ are given by a standard state potential $\mu_i^0$ and due to thermodynamic product inhibition, an adjustment due to concentrations of , i.e., $\mu_i = \mu_i^0 - \log O_i/R_i$.

Each species $\alpha$ may exploit a specific subset of these transformations specified by kinetic coefficients $e_{i\alpha} \geq 0$; e.g., if $e_{i\alpha} \neq 0$, species $\alpha$ can transform $R_i \to O_i$ with kinetic coefficient $e_{i\alpha}$, releasing each electron from transformation $R_i \to O_i$ at a potential $\mu_i$, which are then absorbed by another transformation $O_j \to R_j$ that the species participates in. The net potential difference drop experienced by the electron is the energy available to this species. In practice, the electrons are transferred by an electron carrier (e.g., NADH) in the cell that is at potential $\mu_{\text{carrier},\alpha}$ intermediate to $\mu_i$ and $\mu_j$. We assume there is only one electron carrier pool common to all redox transformations in species $\alpha$. Multiple electron carrier pools would in principle correspond to different carrier potentials, one for each carrier molecule.

We assume that detailed balance is broken across the redox tower because some transformations, say $R_j \to O_j$ are coupled to an external energy (but not matter) source (e.g., coupling the transformation $H_2O$ $\to$ $O_2$ to sunlight during photosynthesis). Consequently, the chemical potential of $R_j \to O_j$ is shifted $\mu_j = \mu_{-j} + \mu_{h\nu}$ where $\mu_j$ is the chemical potential for the reverse transformation $O_j \to R_j$ (not coupled to light). In the simulations in the main text, only one special transformation, corresponding to $j = 3$ ($R_3 \to O_3$) couples to light. This is analogous to the $H_2O$ to $O_2$ transformation. Importantly, the reverse transformation, $O_3 \to R_3$ (analogous to $O_2$ to $H_2O$) does not couple to light and has its adjusted potential given by the standard formula in the previous paragraph.

To specify species and resource dynamics, we first need to compute the per capita fluxes $f_{i\alpha}$ of all transformations $O_i \to R_i$ due to each species $\alpha$. Since multiple microbial species could catalyze the same transformation, and because each transformation cannot proceed faster than a certain timescale, we model $f_{i\alpha}$ using the following expression, similar to Michaelis-Menten kinetics with multiple competing enzymes (here, species):

$$f_{i\alpha} = \frac{\cdot O_i}{(K_N + \sum_{\beta=1}^{S} \frac{N_\beta}{k_{i\beta}})}, \tag{S1}$$

where $k_{i\alpha}$ is the per capita kinetic coefficient corresponding to species $\alpha$ and molecule $i$, $N_\alpha$ is the abundance of species $\alpha$, $O_i$ is the concentration of the corresponding substrate of the transformation, and $K_N$ is a half-saturation constant, chosen arbitrarily to be 1. From thermodynamics, $k_{i\alpha}$ is given by the force-flux relation:

$$k_{i\alpha} = e_{i\alpha} \cdot (1 - e^{-\Delta\mu_{i\alpha}}) \tag{S2}$$

where $\Delta\mu_{i\alpha}$ is the change in potential of an electron released by transformation $O_i \to R_i$ and captured by the electron carrier. Hence:

$$\Delta\mu_{i\alpha} = \mu_{\text{carrier},\alpha} - \left(\mu_i^0 - \log\left(\frac{O_i}{R_i}\right)\right), \tag{S3}$$

where $\mu_i^0$ is the standard state chemical potential of the transformation $O_i \to R_i$ and $\mu_{\text{carrier},\alpha}$ is the chemical potential of the electron carrier for in species $\alpha$. Here, $k_{i\alpha}$ is the net forward rate of the transformation ($J_+/J_-$ in thermodynamics), and $\Delta\mu_{i\alpha}$ is the corresponding energy difference. $e_{i\alpha}$ is a kinetic constant that specifies the per capita per unit concentration rate at which species $\alpha$ can perform transformation $i$, as described above (0 implying $\alpha$ cannot perform the transformation).

Together, all fluxes $f_{i\alpha}$ change the concentrations of $O_i, R_i$ through the following dynamics:

$$\frac{dO_i}{dt} = -\sum_{\alpha=1}^{S} f_{i\alpha} N_\alpha + \sum_{\beta=1}^{S} f_{i\beta} N_\beta \tag{S4}$$

where $f_{i\alpha}$ is the flux of the transformation $O_i \to R_i$ performed by an individual of species $\alpha$ (as given in equation (1)), and $f_{i\beta}$ is the flux of the transformation $R_i \to O_i$ (similar to equation (1), but proportional to the reactant concentration $R_i$, not $O_i$ by individuals of species $\beta$. The first sum goes over all species transforming $O_i \to R_i$ and the second sum over species capable of the reverse. Similar equations hold for $R_i$.

Each species $\alpha$ extracts energy with flux $\mathcal{E}_{\text{tot}}^{\alpha}$ by coupling electrons between transformations at different potentials:

$$\mathcal{E}_{\text{tot}}^{\alpha} = \sum_{i=1}^{2R} \mathcal{E}_{i\alpha} = \sum_{i=1}^{2R} f_{i\alpha} \cdot \Delta\mu_{i\alpha}. \tag{S5}$$

A species grows in abundance if this captured energy exceeds a prescribed per capita maintenance energy $\mathcal{E}_{\text{maint}}$:

$$\frac{1}{N_\alpha}\frac{dN_\alpha}{dt} = \mathcal{E}^\alpha_{\text{tot}} - \mathcal{E}_{\text{maint}}. \tag{S6}$$

Finally, to balance all cycles at steady state, i.e., to conserve matter, as species in the ecosystem couple different half-reactions and transform resources from one form to another, all resource cycles must be balanced (Fig. 1e). Together, these constraints can be summarized as:

$$\underline{\text{electron conservation:}} \qquad \sum_{i=1}^{2R} f_{i\alpha} = 0, \tag{S7}$$

$$\underline{\text{energy requirement:}} \qquad \sum_{i=1}^{2R} f_{i\alpha}\Delta\mu_{i\alpha} = \mathcal{E}_{\text{maint}}, \tag{S8}$$

$$\underline{\text{matter conservation:}} \qquad \sum_{\alpha=1}^{S} N_\alpha f_{i\alpha} = 0. \tag{S9}$$

The last equation amounts to assuming that the ecosystem is fully closed to matter, and open only to an external source of energy (here, light energy $\mu_{h\nu}$) that breaks detailed balance for the chemical potentials $\mu_i$. While we use a fully materially closed ecosystem as an extreme case to illustrate our model, our key results hold for partially closed ecosystems as well (Fig. S3), where some of the resources can be exchanged with the environment and equation (S4) is modified as follows:

$$\frac{dO_i}{dt} = -\sum_{\alpha=1}^{S} f_{i\alpha}N_\alpha + \sum_{\beta=1}^{S} f_{i\beta}N_\beta + \kappa_i - \delta_i O_i, \tag{S10}$$

where $\kappa_i$ specifies the rate of influx of molecule $O_i$, and $\delta_i$ specifies its specific (per unit concentration) dilution rate.

### II. COMPLETE SET OF DYNAMICAL EQUATIONS FOR A 2-SPECIES, 2 CYCLE SYSTEM

For concreteness, we now explicitly write down the set of dynamical equations for the simplest possible ecosystem in our model: a 2-species system (e.g., heterotroph-phototroph) with 2 resource cycles (e.g.,

$H_2O/O_2$ and $CO_2$/org C). This would be an ecosystem where $S = 2$ and $R = 2$, as opposed to ecosystems with $R = 3$ and $S$ between 3 and 10 that we study in the main text. We will assume that the $O_1/R_1$ pair corresponds to $CO_2$/org C and $O_2/R_2$ pair corresponds to $O_2/H_2O$ respectively.

We will represent the phototroph — which transforms $R_2 \to O_2$ and $O_1 \to R_1$ — with label $P$ and abundance $N_P$, and heterotroph — which transforms $O_2 \to R_2$ and $R_1 \to O_1$ — with label $H$ and abundance $N_H$, respectively. Analogous to photosynthesis, only the $R_2 \to O_2$ transformation will coupled to light. Following the previous section, the population dynamics for the two species are given by the following equations:

$$\frac{1}{N_P}\frac{dN_P}{dt} = \frac{R_2}{K_N + \frac{N_P}{e_{2P}(1-\exp(-\Delta\mu_{2P}))}}(\Delta\mu_{2P} - \mu_{h\nu}) + \frac{O_1}{K_N + \frac{N_P}{e_{1P}(1-\exp(-\Delta\mu_{1P}))}}\Delta\mu_{1P} - \mathcal{E}_{maint}, \tag{S11}$$

$$\frac{1}{N_H}\frac{dN_H}{dt} = \frac{O_2}{K_N + \frac{N_H}{e_{2H}(1-\exp(-\Delta\mu_{2H}))}}\Delta\mu_{2H} + \frac{R_1}{K_N + \frac{N_H}{e_{1H}(1-\exp(-\Delta\mu_{1H}))}}\Delta\mu_{1H} - \mathcal{E}_{maint}, \tag{S12}$$

where:

$$\Delta\mu_{i\alpha} = \mu_{\text{carrier},\alpha} - \left(\mu_i^0 - \log\left(\frac{O_i}{R_i}\right)\right) \tag{S13}$$

for all $i \in 1, 2$ and $\alpha \in P, H$. The first term in equation (S11) has an additional term $\mu_{h_\nu}$ that indicates that the transformation $R_2 \to O_2$ is coupled to light and breaks detailed balance. The dynamics for the $\mu_{\text{carrier},\alpha}$ are given by the following equations:

$$\frac{d}{dt}\mu_{\text{carrier},P} = \frac{R_2}{K_N + \frac{N_P}{e_{2P}(1-\exp(-\Delta\mu_{2P}))}} - \frac{O_1}{K_N + \frac{N_P}{e_{1P}(1-\exp(-\Delta\mu_{1P}))}}, \tag{S14}$$

$$\frac{d}{dt}\mu_{\text{carrier},H} = \frac{O_2}{K_N + \frac{N_H}{e_{2H}(1-\exp(-\Delta\mu_{2H}))}} - \frac{R_1}{K_N + \frac{N_H}{e_{1H}(1-\exp(-\Delta\mu_{1H}))}}. \tag{S15}$$

The nutrient dynamics are given by the following dynamical equations:

$$\frac{dR_1}{dt} = -\frac{R_1}{K_N + \frac{N_H}{e_{1H}(1-\exp(-\Delta\mu_{1H}))}}N_H + \frac{O_1}{K_N + \frac{N_P}{e_{1P}(1-\exp(-\Delta\mu_{1P}))}}N_P, \tag{S16}$$

$$\frac{dO_1}{dt} = \frac{R_1}{K_N + \frac{N_H}{e_{1H}(1-\exp(-\Delta\mu_{1H}))}}N_H - \frac{O_1}{K_N + \frac{N_P}{e_{1P}(1-\exp(-\Delta\mu_{1P}))}}N_P, \tag{S17}$$

$$\frac{dR_2}{dt} = -\frac{R_2}{K_N + \frac{N_P}{e_{2P}(1-\exp(-\Delta\mu_{2P}))}}N_P + \frac{O_2}{K_N + \frac{N_H}{e_{2H}(1-\exp(-\Delta\mu_{2H}))}}N_H, \tag{S18}$$

$$\frac{dR_2}{dt} = +\frac{R_2}{K_N + \frac{N_P}{e_{2P}(1-\exp(-\Delta\mu_{2P}))}}N_P - \frac{O_2}{K_N + \frac{N_H}{e_{2H}(1-\exp(-\Delta\mu_{2H}))}}N_H. \tag{S19}$$

Examples of dynamics from this system are shown in Fig. S1. Similar examples from more complex many species ecosystems with $R = 3$ are shown in Fig. S2.

### III. PARAMETERS AND INITIAL CONDITIONS USED FOR SIMULATIONS

| Symbol | Interpretation | Typical value (if applicable) |
|---|---|---|
| $R$ | Number of resource cycles | 3 |
| $S$ | Number of species added | 3–10 |
| $N_{pool}$ | Number of species in the random ensemble | 1,000 |
| $K_N$ | Half-saturation coefficient for flux | 1 |
| $O_i + R_i$ | Total amount for resources in each cycle | 1 |
| $e_{i\alpha}$ | Kinetic coefficient for transformation $i$ by species $\alpha$ | $\mathcal{U}(0,1)$ |
| $\mu_i^0$ | Standard state potentials of redox pairs | $\{+0.8, -0.4, -0.1\}$ |
| $\mu_{h\nu}$ | Potential of external driving force (light) | 2 |

For each simulation, we assumed random initial conditions for all species abundances and electron carrier potentials, such that they were uniformly distributed between 0 and 1. For each resource cycle $i$, we uniformly partitioned the total resource amount $O_i + R_i$ at random. We ran all simulations using the Radau solver for 1,000 time steps. We simulated equations for species abundances, resource concentrations and electron carrier potentials to steady state (equations (S7)–(S9)).

As $O_i, R_i$ concentrations and their corresponding chemical potentials changed, each species reorganized its internal fluxes to balance the net flow of electrons and subsequently captured a different energy flux from the environment. Some species continued to capture more than $\mathcal{E}_{\text{maint}}$ and grow, while others captured less than $\mathcal{E}_{\text{maint}}$ and decreased in abundance, some even going extinct. This feedback between species and resources continued until the ecosystem self-organized into a steady state where individuals of every surviving species captured energy flux $\mathcal{E}_{\text{maint}}$ with a balanced internal redox electron flux, and all resource cycles were balanced at certain fluxes $\phi_i$. Together, all surviving individuals in the ecosystem captured a total energy flux $\mathcal{E}_{\text{tot}}^{\text{eco}} = \sum_{\alpha=1}^{S} \mathcal{E}_{\text{tot}}^{\alpha} N_\alpha$ (Fig. 2a). By simulating 1,000 such ecosystems, we sampled a large space of ecosystems that obeyed thermodynamic constraints arising from using redox transformations as resources.

### IV. MEASURING THE VOLUMES OF ECOSYSTEM SOLUTION SPACES

To compare the volumes of the space of possible ecosystem solutions — both in terms of species abundances and nutrient cycle fluxes — simulated using our model, we projected all solutions to a common, two-dimensional space. This projection was necessary for any comparison because the species and flux spaces have different inherent dimensionalities ($S = N_{pool}$ and $R = 3$ respectively). To preserve pairwise Euclidean distances between ecosystems and allow for a fair comparison of projected volumes, we first normalized all Euclidean distances in species and flux spaces by the maximum Euclidean distance, and then used multi dimensional scaling (MDS), for which standard algorithms exist. We projected all species abundance and cycle flux vectors to a common two-dimensional space (Fig. 2d). To estimate the volume of each space, we measured the 'volume' (area) of the convex hull of the projected set of species and flux points respectively. We then repeated this procedure for each value of $S$, each of which we had an ensemble of 1,000 ecosystems (points) for. We then normalized all measured volumes by the the largest volume (so that the largest volume was 1), and plotted this for the species (green) and fluxes (red) solutions in Fig. 2e.

### V. BUILDING ENSEMBLES OF MACHINE ECOSYSTEMS

To build ensembles of ecosystems of machines, we performed constrained optimization for equations (S7) and (S9), instead of dynamically evolving the equations (S4), (S6) and equations of the form (S14-15). Specifically, we used the same set of parameters as for biological ecosystems, but for each machine ecosystem with randomized abundances, we numerically found a solution to the constraints (S7) and (S9) corresponding to them for a randomized set of species abundances, chosen uniformly in log-space between $10^{-5}$

and 1. To find each solution, we used standard non-linear least squares methods implemented in SciPy. For ecosystems of machines with optimized abundances for the same parameters, we chose species abundances so that they maximized the total energy extraction $\mathcal{E}_{\text{tot}}^{\text{eco}} = \sum_{\alpha=1}^{S} \mathcal{E}_{\text{tot}}^{\alpha} N_\alpha$, while subject to the constraints (S7) and (S9) corresponding to the parameters. For this, we used the SciPy constrained optimization routine 'minimize' with method SLSQP.

### VI. SUPPLEMENTARY FIGURES

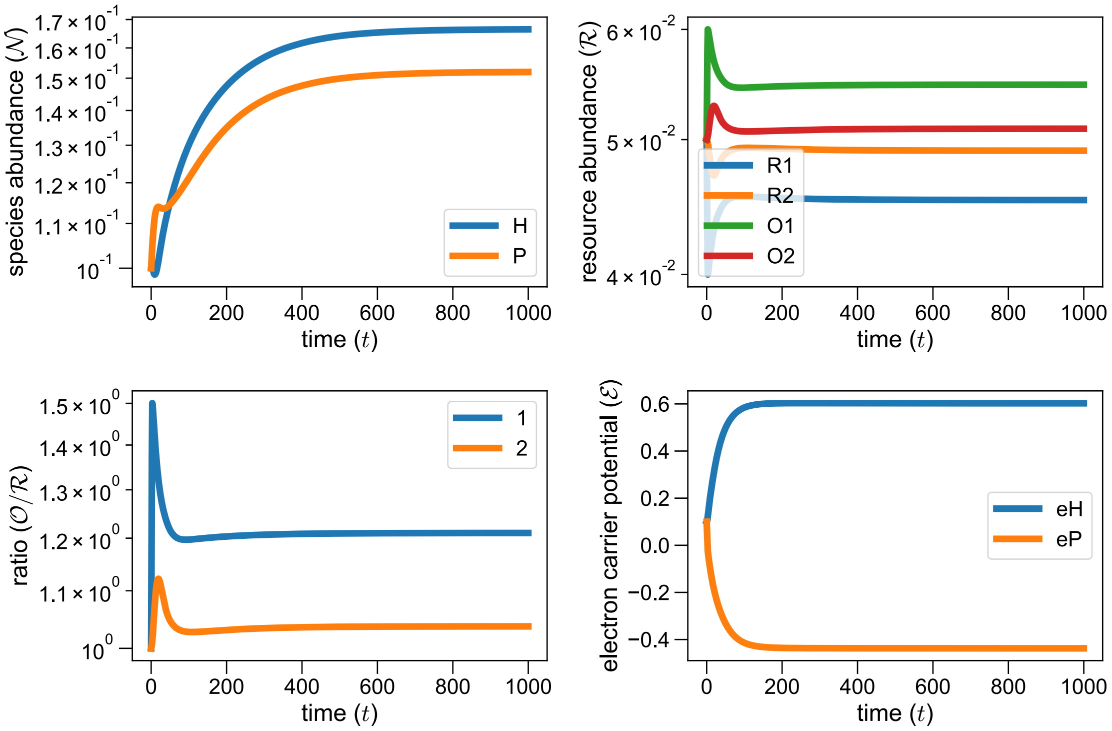


FIG. S1. **Example of dynamics in a 2-species, 2-resource cycle ecosystem.** Example of dynamics of an ecosystem with $S = 2$ and $R = 2$, corresponding to the phototroph-heterotroph system outlined in section II, with default parameters and random initial conditions (section III). The top-left panel shows the species dynamics ($H$ representing the heterotroph in blue and $P$, the phototroph in orange). The top-right panel shows the resource dynamics (legend shown), while the bottom panels show the dynamics of resource ratios (left) and electron carrier potentials in both species (right).

.

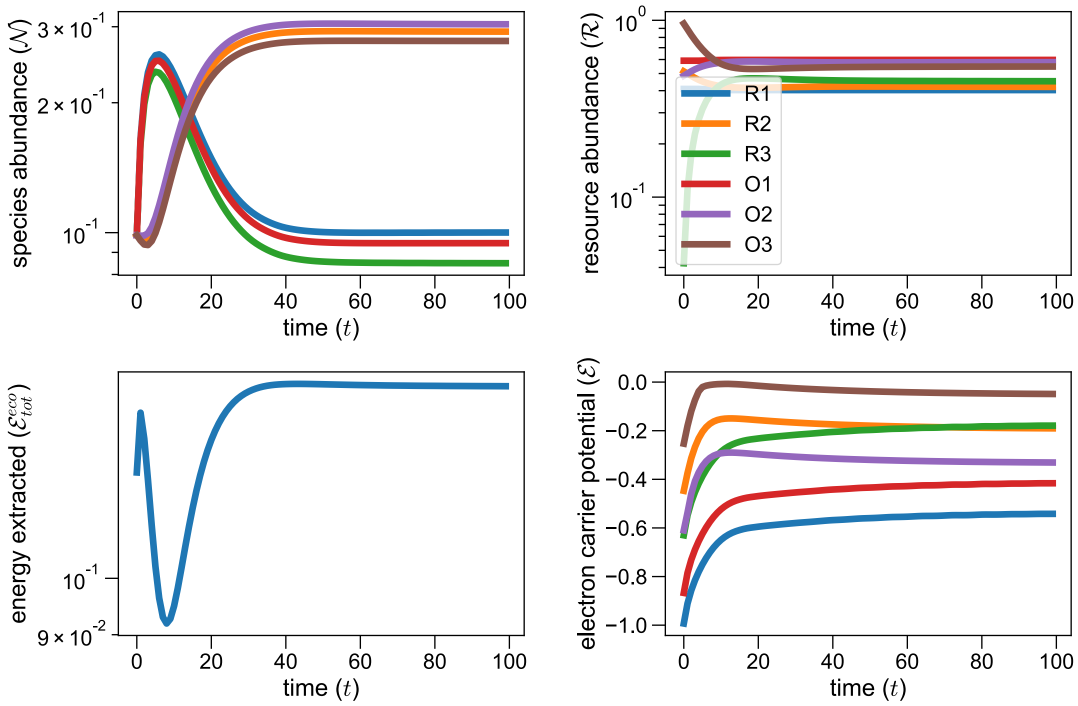


FIG. S2. **Example of dynamics in a 6-species, 3-resource cycle ecosystem.** Example of dynamics of an ecosystem with $S = 6$ and $R = 3$, corresponding to a complex ecosystem outlined in the main text, Fig. 2, with default parameters and random initial conditions (section III). The top-left panel shows the species dynamics (each color representing a different random species). The top-right panel shows the resource dynamics (legend shown), while the bottom panels show the dynamics of the total energy extracted by the ecosystem (left) and electron carrier potentials in all species (right).

.

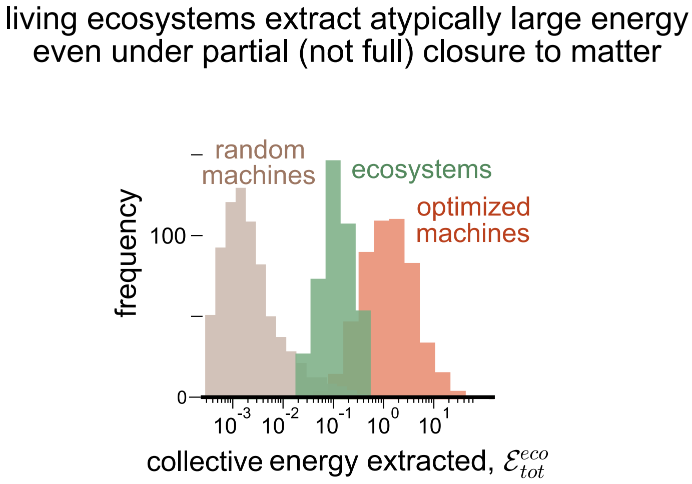


FIG. S3. **Our main result is robust even if ecosystems are only partial closed to matter.** Histograms of the total energy extracted $\mathcal{E}^{\rm eco}_{\rm tot}$ by ecosystems of machines with random abundances (gray; representing initial conditions of ecosystem assembly), ecosystems of self-replicators with abundances self-organized by birth-death dynamics based on maintenance energy (green), and machines with abundances chosen to maximize $\mathcal{E}^{\rm eco}_{\rm tot}$ (orange). Similar to Fig. 4b, but simulated using one of the 3 resource cycles ($O2 \leftrightarrow R_2$) open to matter ($\kappa_2 = 0.1, \delta_2 = 0.1$). On average, ecosystems still extract energy closer to optimal than those comprising machines with randomized abundances.

.

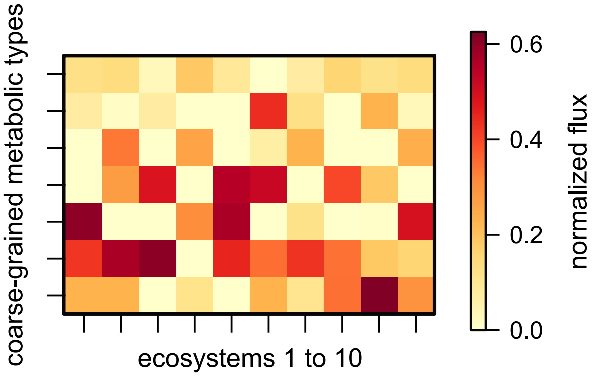


FIG. S4. **Ecosystem structure is more convergent when species are coarse-grained by metabolic types.** Heatmaps showing examples from 10 of the 1,000 randomly assembled ecosystems in Fig. 2, showing the combined abundances of species when grouped by their "metabolic type", i.e., by the subset of transformations they can perform with $e_{i\alpha} \neq 0$. Each row shows a metabolic type, while each column shows an example ecosystem.

.

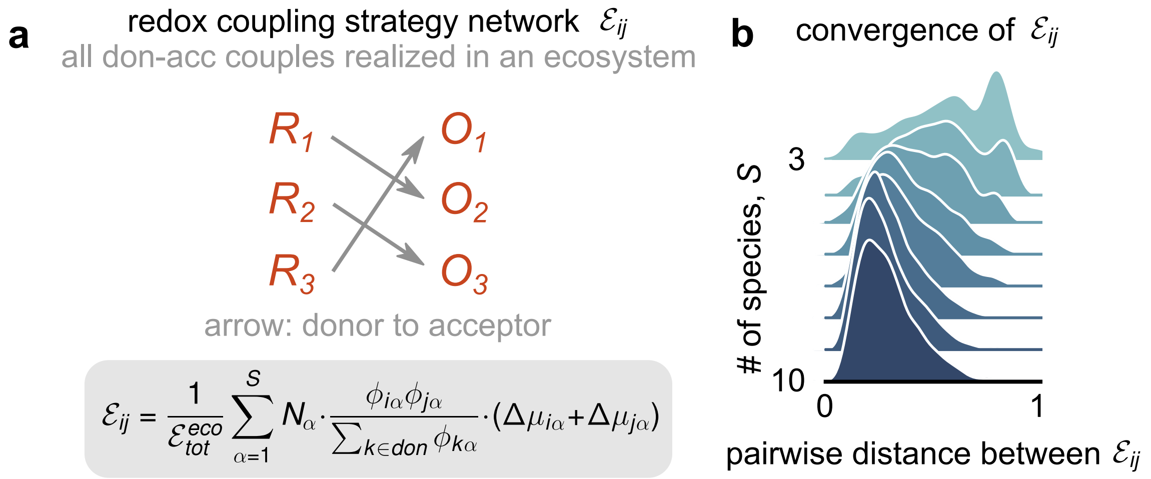


FIG. S5. **Global redox "strategies" converge with increasing species diversity.** (a) Schematic showing an example of a global redox strategy for an entire ecosystem, which quantifies the fraction $\mathcal{E}_{ij}$ of the total ecosystem energy flux $\mathcal{E}^{eco}_{tot}$ that is obtained by coupling transformations $i$ and $j$. We compute $\mathcal{E}_{ij}$ as shown in the gray box, where $\phi_{i\alpha} = f_{i\alpha}N_\alpha$ is the contribution to the total flux in resource $i$ by all individuals of species $\alpha$. Notably, $\sum_{i\neq j}\mathcal{E}_{ij} = 1$ is a matrix that is normalized by definition. (b) Histograms of the pairwise Euclidean distance between the global redox strategies $\mathcal{E}_{ij}$, as a function of number of species added $S$, for all ecosystems simulated in Fig. 2. The average distance (and its variance) decreases with increasing diversity, suggesting that the strategies implemented across different ecosystems become more similar (converge).

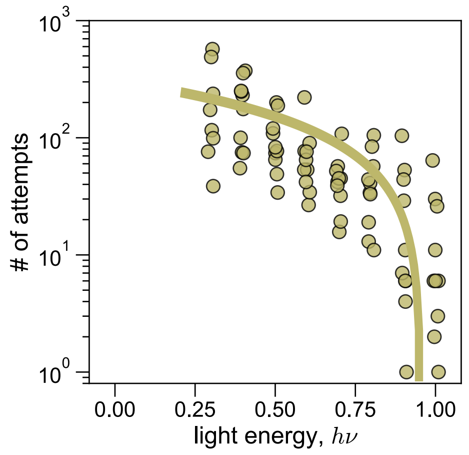


FIG. S6. **Once enough driving light energy is available, randomly assembled ecosystems rarely collapse.** Scatter plot showing the number of simulations with random initial conditions (section III) that we need to run before we arrive at an ecosystem that can self-organize to capture nonzero energy, i.e., that does not collapse. We show the number of attempts as a function of the light energy driving the ecosystems $\mu_{h\nu}$, as explained in section I. Ecosystems cannot self-organize below a minimum $\mu_{h\nu}$ as in Fig. 3. Once the driving energy is large enough, ecosystems can almost always self-organize (the number of attempts become close to 1).

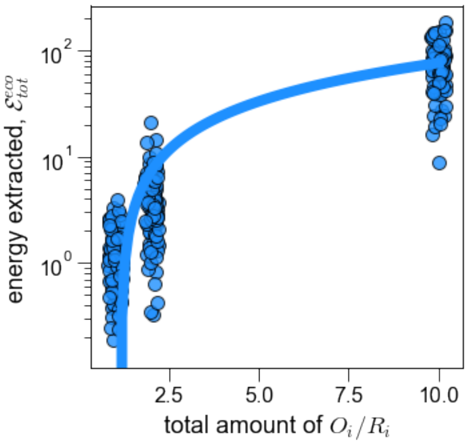


FIG. S7. **Once there is sufficiently large light energy, the total amount of resources limit the energy extracted by ecosystems.** Scatter plot showing the total energy extracted $\mathcal{E}_{tot}^{eco}$ as a function of the total amount of resources $\sum(O_i + R_i)$ supplied to closed ecosystems. Each point represents a randomly assembled ecosystem from a large ensemble of species, as in Figs. 2 and 3, but assembled with different total resource amounts.

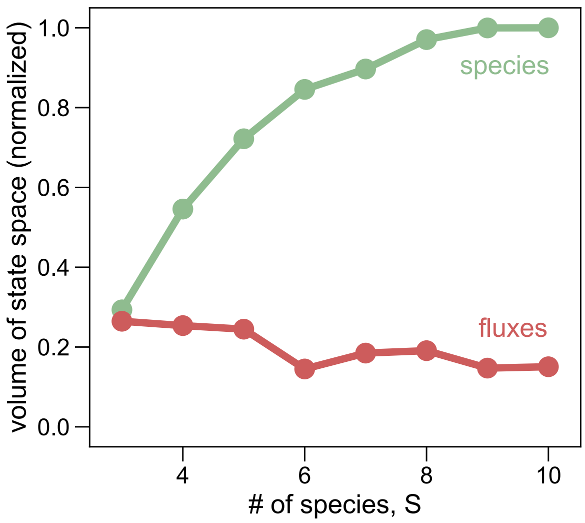


FIG. S8. **Our results about flux convergence are robust to the addition of more resource cycles.** Line plot, similar to Fig. 2e, but with $R = 5$ instead of 3. The plot shows how the volume of the species (green) and flux (red) spaces scales with the number of species added, $S$, in assembled ecosystems. As in Fig. 2e, the flux space volume grows much slower than species space volume, indicating convergence in the function (fluxes) of self-organized ecosystems.

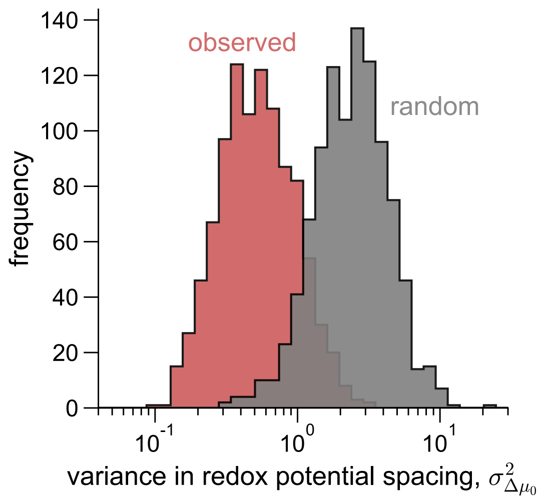


FIG. S9. **Self-organized redox potentials are more equally spaced than expected by chance.** Histograms of the variance in spacing between adjusted redox potentials for ecosystems at steady state (red), compared with the variance in spacing when the potentials are randomized, i.e., spread uniformly in the same range. The observed potentials show roughly 10-fold lower variance in spacing.
